# Mapping quantitative trait loci underlying circadian light sensitivity in *Drosophila*

**DOI:** 10.1101/135129

**Authors:** Adeolu B. Adewoye, Sergey V. Nuzhdin, Eran Tauber

**Author notes:** Corresponding author: E. Tauber. Present address: Wolfson School of Mechanical and Manufacturing Engineering, Centre for Biological Engineering, Loughborough University Loughborough, United Kingdom (Adeolu B. Adewoye).

## Abstract

Despite the significant advance in our understanding of the molecular basis of light entrainment of the circadian clock in *Drosophila*, the underlying genetic architecture is still largely unknown. The aim of this study was to identify loci associated with variation in circadian photosensitivity, which are important for the evolution of this trait. We have used complementary approaches that combined quantitative trait loci (QTL) mapping, complementation testing and transcriptome profiling to dissect this variation.

We identified a major QTL on chromosome 2, which was subsequently fine-mapped using deficiency complementation mapping into two smaller regions spanning 139 genes, some of which are known to be involved in functions which have been previously implicated in light entrainment. Two genes implicated with the clock and located within that interval, *timeless* and *cycle*, failed to complement the QTL, indicating that alleles of these genes contribute to the variation in light response. Specifically, we find that the *timeless* s/*ls* polymorphism that has been previously shown to constitute a latitudinal cline in Europe, is also segregating in our recombinant inbred lines, and is contributing to the phenotypic variation in light sensitivity.

We have also profiled gene expression in two recombinant inbred strains that differ significantly in their photosensitivity, and identified a total of 368 transcripts that showed differential expression (FDR < 0.1). Out of 131 transcripts that showed a significant RIL by treatment interaction (i.e. putative expression QTL), four are located within QTL2

## Introduction

How genetic variation allows organisms to adapt to their changing environments, either temporally or spatially, is a major question in evolutionary genomics. For most organisms, light is the primary environmental cue that regulates daily rhythms, by aligning the circadian clock to the 24-h light/dark cycle (LeGates *et al.* 2014). In the laboratory, light entrainment of the clock is often studied by light-pulse experiments, where brief light stimulus is presented to subjects maintained in continuous darkness (Daan and Aschoff. 2001). Most organisms, including *Drosophila*, respond to the stimulus by shifting their free-running rhythm to a new phase. The amount of phase shift, depicted as a phase-response curve (PRC), depends on the timing (phase), intensity and duration of the stimulus, and the light-sensitivity of the subject (Johnson *et al*. 2003). Many species share the same basic PRC, with early night stimuli causing phase delays, and late night stimuli leading to phase advances, although the precise shape may vary between species (Johnson *et al*. 2003), or populations within a species (Pittendrigh *et al*. 1991).

Extensive research over the past five decades has elucidated the molecular basis of the circadian clock in *Drosophila* (Dubowy and Sehgal. 2017). Briefly, clock proteins PERIOD (PER) and TIMELESS (TIM) negatively regulate their own transcription via interaction with the transcription activators CLOCK (CLK) and CYCLE (CYC). This negative feedback loop drives a cyclical expression pattern where accumulation of protein levels of PER and TIM is associated with reduced level of the mRNA (Pegoraro and Tauber. 2011). The proteins PDP1 and Vrille form a second feedback loop that drive the oscillation of *Clk* mRNA (Cyran *et al*. 2003; Hardin *et al*. 2003; Zhou *et al*. 2016). The proteins Clockwork Orange (CWO) is another repressor of the CLK-CYC complex (Kadener *et al.* 2007). The transcriptional-translation feedback loops are enhanced by post-translational mechanisms, particularly phosphorylation. This is carried out by kinases such as DOUBLETIME (DBT), Shaggy (SGG) and NEMO (Kloss *et al*. 1998; Price *et al*. 1998), and by phosphatases such as Protein Phosphatase 2A (Sathyanarayanan *et al*. 2004) and Protein Phosphatase 1 (Fang *et al.* 2007).

While many of the core components of the circadian clock have been identified, the extent of natural genetic variation associated with the circadian clock is largely unknown. This variation is expected to be substantial, reflecting different molecular adaptations that have evolved in wild populations in response to various environmental cues. For example, there is evidence for molecular variations in clock genes that exhibit geographical clines, following the latitudinal change of photoperiod and temperature (Kyriacou *et al*. 2008). Identifying and characterising these variations is essential for understanding of the clock and its evolution.

An early study (Pittendrigh *et al.* 1991) has shown that light sensitivity in natural populations of *Drosophila auraria* follows a latitudinal cline with an apparently reduced light response in northern populations. This was explained as an adaptive response to the extremely long days of the summer. Although the genetic basis for this phenotypic cline in *D. auraria* is still unknown, this observation indicated that genetic variation underlying this trait is present in wild populations.

Later, a natural polymorphism in the light-sensitive clock protein TIM was identified in *D. melanogaster* (Tauber *et al*. 2007). A single-base insertion/deletion, situated between two alternative translation starts, results in two alleles, *ls-tim* allele (the insertion) produces both long and short isoforms of the protein whilst the *s-tim* allele (the deletion) generates only the shorter isoform. This polymorphism follows a robust latitudinal cline and is maintained by directional selection. Light pulse experiments have indicated that *ls-tim* flies are less light responsive than flies carrying *s-tim* (Sandrelli *et al.* 2007).

Quantitative trait loci (QTL) mapping is a popular approach for identifying loci that underlie phenotypic variation of a complex trait. Attempts to map circadian traits by QTL mapping have been previously made in plants (Darrah *et al.* 2006; Michael and McClung. 2003), mice (Yoshimura *et al.* 2002a) and fungi (Kim *et al.* 2007), but not in *Drosophila.* The available powerful tools for fly genetics, and its compact genome, offer a rapid means to move from an identified QTL to a specific gene and the underlying polymorphism (Mackay. 2001). The current study was aimed at identifying QTLs for circadian photo-sensitivity and isolating the underlying candidate genes.

## Materials and methods

### Mapping population

A set of 123 recombinant inbred lines (RIL) were randomly selected from a larger population of 300 RIL previously described (Bergland *et al.* 2008). This panel was generated from two isofemale lines from a wild population in Winters, California collected in 2001. These parental lines were made isogenic by inbreeding and were used for generating 500 isogenic lines, which were crossed with each other for 15 generations. This was followed by full sib crossing for 15 generations resulting in the RILs panel.

The lines were maintained in vials of yeast-sucrose-agar medium under (∼70 % relative humidity, at either 18°C or 25 °C and 12h: 12h light cycle).

### Light pulse experiment

The locomotor activity of the flies was measured using the DAM2 *Drosophila* monitors (Trikinetics Inc, Waltham, USA). Single flies were placed in glass tubes (10cm x 0.5 cm) that were filled with 2 cm sugar/agar medium. The monitors were placed in an incubator at 25°C, ∼70% humidity. The flies were entrained to a light-dark cycle (LD 12:12) for 4 days and then allowed to free-run for 3 days in constant darkness (DD). This entrainment regime (4 days LD, 3 days DD) was repeated with the same set of flies in the following week, but with a 20 min light pulse at ZT15 at in the last dark phase of the LD cycle. The light pulse (1500 lux) was delivered by two fluorescent lamps (Philips MCFE 20W/35) mounted behind a diffuser.

For each fly, the phase difference between the reference phase (second day DD, off-set of locomotor activity bout) and the off-set of the second day in DD after the 20-min light pulse was taken as the phase delay. Note that by the second day, all flies complete their phase shift (Suri *et al.* 1998). The reference and the response phase were determined manually by visual inspection of each fly activity profile.

### QTL analysis

The broad sense heritability (*h*^2^) for the RIL population, and the genome scans for QTL were performed using composite interval mapping (CIM) (Zeng *et al*. 1999) and multiple interval mapping (MIM) in Windows QTL Cartographer version 2.5 program (WinQTLCart) (Wang *et al.* 2007). The threshold value for the CIM analysis were determined at 5% level of 1000 permutations (Churchill and Doerge. 1994). The confidence intervals (95%) for the QTL were determined by identifying the region in which the LOD score is within 1.5 of its highest peak, and markers spanning the QTL as determined from the map were used. The cytological positions and the putative genes were estimated using the markers’ physical location from Ensembl release 56 and the FlyMine database (Lyne. *et al*. 2007)

### Complementation tests

A total of eleven deficiency stocks spanning QTL 2 (22B1 – 25B1) and QTL 3 (71E1 – 79A1) (obtained from Bloomington Drosophila Stock Centre, see Supplementary Table S1) were used to fine-map these QTL intervals. Two RI lines: RIL104 and RIL58, which showed high and low light response respectively, were chosen for further analysis of the QTL as the parental lines were not available. In addition to their difference in light response, these lines also showed a close resemblance to the parental genotypes along the QTL regions. Males from RIL58 and RIL104 were crossed with virgin females from each of the deficiency lines, resulting in four F1 genotypes: *58/Df, 58/Bal, 104/Df* and *104/Bal*, where *Df* and *Bal* refer to deficiency and the balancer chromosomes respectively. A two-way ANOVA was carried out using the RIL (L) and the genetic background (G), which were treated as fixed effects. Quantitative failure to complement the QTL alleles is inferred when the interaction of the two factors (L × G) is significant. Specifically, we required that difference in mean light response between the genotypes *58/Df 104/Df* would be significantly greater than that between *58/Bal* 104*/Bal.*

Complementation tests were also carried out using: *y*^*1*^ *w*; *Thor*^*2*^, *w*; *tim*^*01*^, *w;cyc*^*01*^ *and clk*^*JRK*^ strains (all within the QTL). Crosses were performed as described above but using the mutant strain and *w*^*1118*^ instead of the Df and the balancer chromosome. It should be noted that in both cases, the genetic background is not identical across the four genotypes that were compared in each test. For example, the *58/Bal* and *58/Df* differ in their overall background, so phenotypic differences may due to loci outside the deletion. Thus, a statistical significant interaction could also be due to various epistatic interactions involving other loci (Service. 2004), and may lead to false positives.

### Sample collection for transcriptomics

Five day post-eclosion males from RIL58 and RIL104 were entrained to four LD cycles. Flies were then split into two groups, one presented with a light pulse for 30 minutes at ZT 15 and a control group kept in the same conditions without the pulse. Flies were collected at ZT 16.5 and flash frozen in liquid nitrogen under red light. Two independent replicates were prepared per condition, totalling 8 samples: 2x lines (L) x treatments (T) x 2 replicates. Total RNA was isolated from fly heads using Invitrogen TRIzol reagent in accordance with the company protocol. RNA concentration was determined using NanoDrop 2000 (Thermo Scientific) and sample quality was assessed by the Agilent 2100 Bioanalyzer (Agilent Technologies, Santa Clara, CA, USA) according to the manufacturers’ protocols.

### cDNA samples processing

Invitrogen superscript plus indirect cDNA kit was used to generate fluorescently labelled cDNA from 35 µg total RNA according to the manufacturer’s protocol (Invitrogen). The purification and quantification were done as recommended by the Invitrogen protocol. The purified Alexa 555 and Alexa 647 labelled cDNA were hybridised to *Drosophila* Oligo 14Kv1 array (Canadian Drosophila Microarray Centre) according to University Health Network (UHN) Microarray Centre amino-allyl (indirect) labelling protocol. The array slides were scanned on Molecular Devices’ GENEPIX 4000 microarray scanner.

### Data processing and statistical analysis

The data was normalized and processed by the *limma* package (Ritchie *et al*. 2015) using the R software (R Development Core Team. 2010). The data were corrected for background intensities and normalised across each slide using the print-tip loess method (Yang *et al.* 2001). Differential expression was assessed by using a linear model and Bayesian fit that generated the moderated t-statistics (Smyth. 2004). The analysis implemented the pooled correlation method to make full use of the duplicate spots on the slide (Smyth *et al.* 2005). Because our main focus are components of the central pacemaker, the data were filtered to include only genes that are expressed in the brain (FlyAtlas mRNA signal > 10). We also filter transcripts that show a fold change of log2FC >= 0.1. The p-values were adjusted using the Benjamini and Hochberg false discovery (FDR) method (Benjamini and Hochberg. 1995). Analysis of enriched gene ontologies (GO terms) among differential expressed genes was carried out using the FlyMine Database (Lyne *et al*. 2007). To carry out analysis of regulatory motifs enrichment, a 2kb upstream sequence was extracted from each candidate gene using BioMart, a data mining tool from Ensembl (Aken *et al.* 2017). We used the AME algorithms (McLeay and Bailey. 2010), implemented in the MEME suite (Bailey *et al*. 2009) to search and identify enriched motifs.

## Results

The RI lines exhibited extensive variation in the magnitude of the phase response to the light pulse, ranging from -1.9 to -5.5 hr delays (Fig. 1), and the difference between the strains was statistically significant (Kruskal-Wallis, *H* (123) =288.18, p=0.001). The estimation of the broad-sense heritability (*h^2^*) for the RIL population was 0.3, suggesting a moderate genetic component for light response in this population.

**Figure 1.**
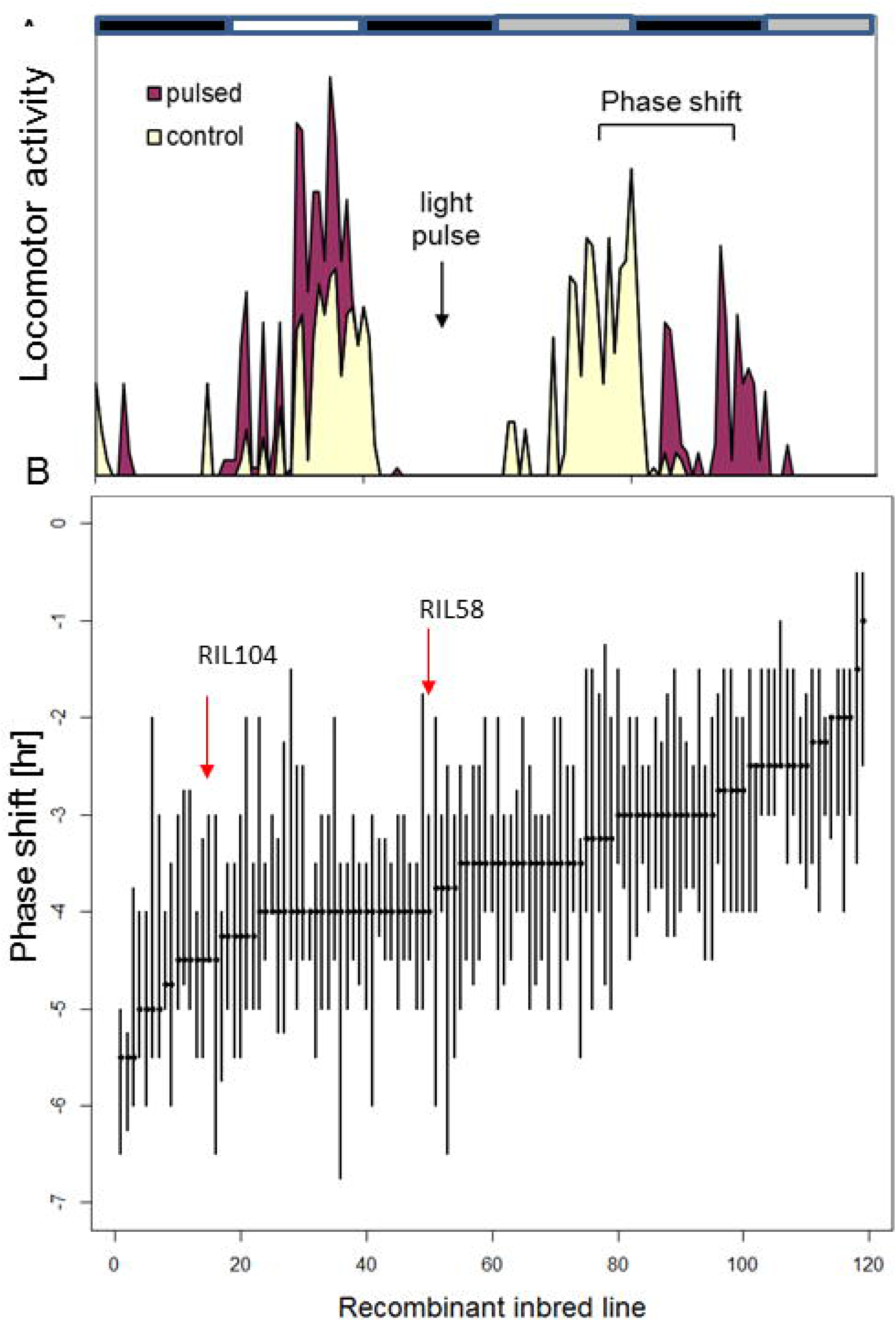
Variation in circadian photosensitivity among RI lines. **A**. diagram showing two days of locomotor activity rhythms for a population of flies that were treated by a light pulse at ZT15 (pulsed) and a non-treated control group. Upon light pulse, the subsequent activity is delayed. The delay interval (bracket), represent the phase delay. **B**. The phase delay response to early light pulse is shown for each of the 123 RI lines. The median and 25-75 percentiles are plotted (20-30 males each line).

We used composite interval mapping (CIM) to identify QTL underlying circadian photosensitivity in the RI lines. One significant QTL was detected with a LOD score of 2.4 (Fig. 2) and three additional suggestive QTLs were indicated by multiple-interval mapping (MIM) (Fig 2, Supplementary Table S2). Together, these QTL explained 30% of the variation in delay response, with the major QTL (QTL2) explaining 12% of the total phenotypic variation observed for this trait. This QTL is located on the left arm of second chromosome (22C1 - 25A3) and spans a genomic region of about 2,600 kb encompassing 316 genes. The QTL on the X chromosome (cytological bands 10C7 -11B12) explains 7.1 % of the overall variation. The other two QTLs are located on the same arm (L) of the third chromosome (72A1 - 78D4 and 62D4 – 66D12) and account for 4.7% and 6.6% of the variation respectively. There was no evidence for any epistatic effect.

**Figure 2.**
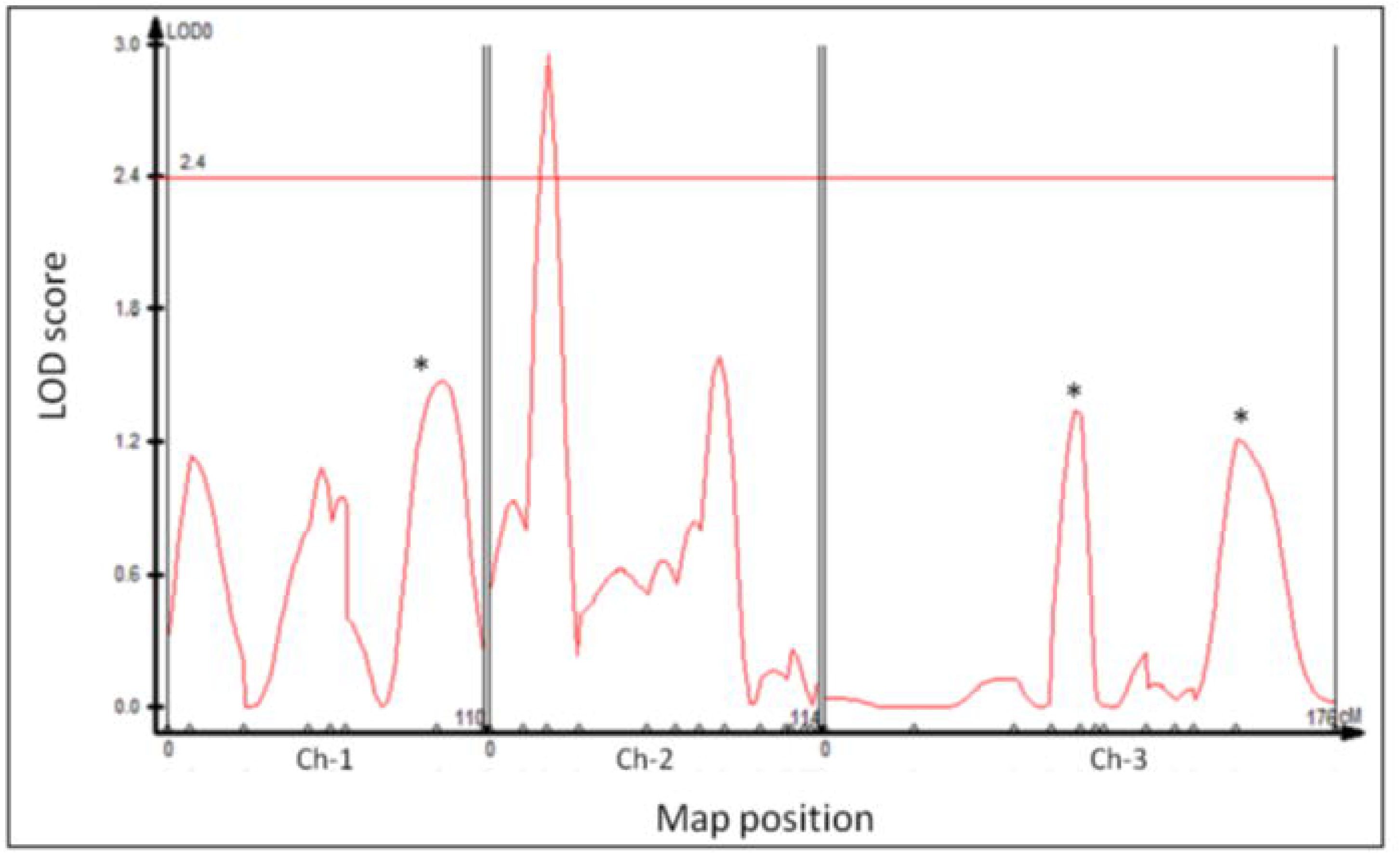
Composite interval mapping of QTL for circadian light sensitivity. Log of odds (LOD) scores and significant thresholds plotted against chromosome location for phase shift response. The horizontal line represents the threshold level (p=0.05) determined by 1000 permutation tests. Marker position is shown on the X-axis. The vertical lines separate chromosomes. Asterisks indicate additional QTL that were suggested by MIM.

Out of the 10 deficiencies that were tested, two failed to complement the QTL alleles (Table S3). These were *Df3133* (22A3 – 22E1) in QTL2 and *Df6411* (74D3-75B5) in QTL3 region (Fig. 3). Additional quantitative complementation tests were performed using mutant strains of three known clock genes, *tim, cyc and Clk* that resides in the QTL. A mutant for the gene *Thor*, which has also been recently implicated in clock function (Nagoshi *et al.* 2010) was also tested. Two of these mutants: *tim*^*0*^ and *cyc*^0^, failed to complement the QTL phenotype, while no significant interaction was found for *Clk*^*jrk*^ and *Thor*^*2*^ (Fig. 3).

**Figure 3.**
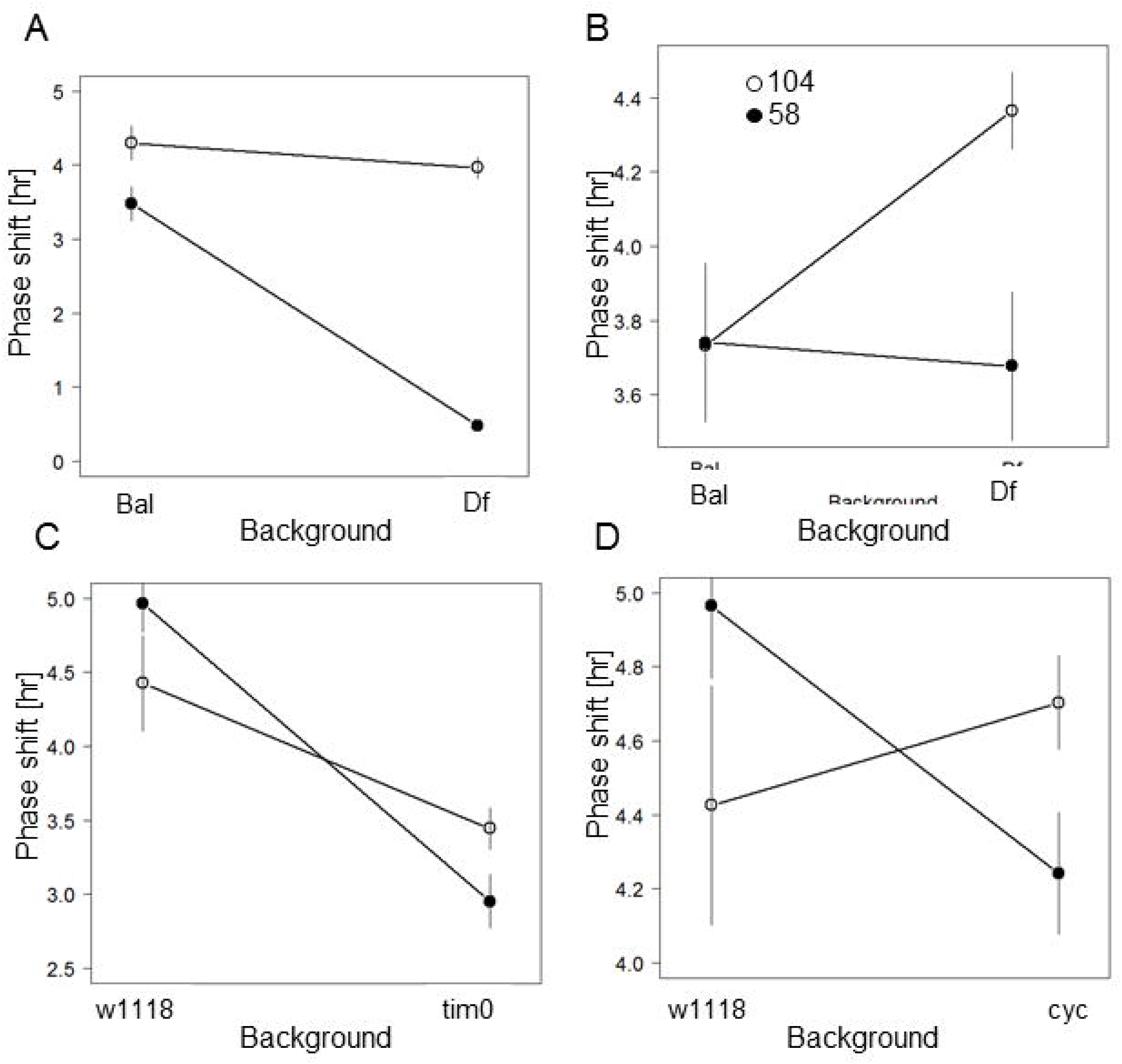
Quantitative complementation to deficiency strains. Interaction plots are shown for RIL58 (filled circles) and RIL104 (open circles) crossed with deficiency strains (**A**) 6411 and (**B**) 3133, which showed a significant Genotype x Line interaction effect (see text). At the bottom are shown complementation test using (**C**) tim^0^ and (**D**) cyc^0^ null mutants. Error bars represent the standard error of the mean. Sample size of each genotype ranged 18 – 32 flies.

Since the complementation tests suggested that variation in *tim* contributes to the major QTL2, we tested whether the previously published *ls-tim* polymorphism segregates in the RIL by genotyping 66 randomly selected lines. Supplementary figure S1 shows that both alleles segregate in the RI strains and that *s-tim* flies responded to light with significantly longer delays than *ls-tim* flies (median delays of -4.0 vs -3.3, Mann-Whitney *U* =*267, df=64, p < 0.0001,*). The RIL104 flies with the longer delay carry *s-tim* while RIL58 (less light sensitive) was *ls-tim,* consistent with the known photosensitivity of these alleles (Sandrelli *et al.* 2007).

In another set of experiments, we used microarrays to examine gene expression of RIL58 and RIL104 following a light pulse at ZT15. Variation in transcriptional response to light between these two lines may represent expression QTL (eQTL), which could be either cis-eQTL (QTL mapped to the location of the expressed gene) or tran-eQTL (QTL is mapped far from the expressed gene).

We profiled gene expression at ZT16.5 as at this time point has been previously used for capturing early expression response to light stimulus (Adewoye *et al.* 2015). The current experiment also revealed an extensive differential expression in both lines, comparing flies that received a light pulse to flies that did not (Fig. 4). At the level of FDR < 0.1, there were 184 differentially expressed genes (DEG) in RIL58, and 284 DEG in RIL104 (Supplementary Table S4). There was substantial difference between the GO terms that were enriched by each of the strains: the top terms in the RIL58 list included regulation of cell cycle (*p* < 0.001), cellular component assembly, as well as protein localisation and cytoskeleton-dependent intracellular transport (Supplementary Table S5). In RIL104 top GO terms were associated with metabolic processes involving galactose, lipids and other metabolites. There were 42 overlapping DEG between the two RILs (Supplementary Table S6), none of which has been previously implicated in light response of the clock. GO analysis of these overlapping genes represented a broad range of biological processes including metal ion binding (*CG10889, msl-2, cg5196*), voltage-gated ionic channel activity *(shaw*) and ATP binding (*pll, sun, grp*).

**Figure 4.**
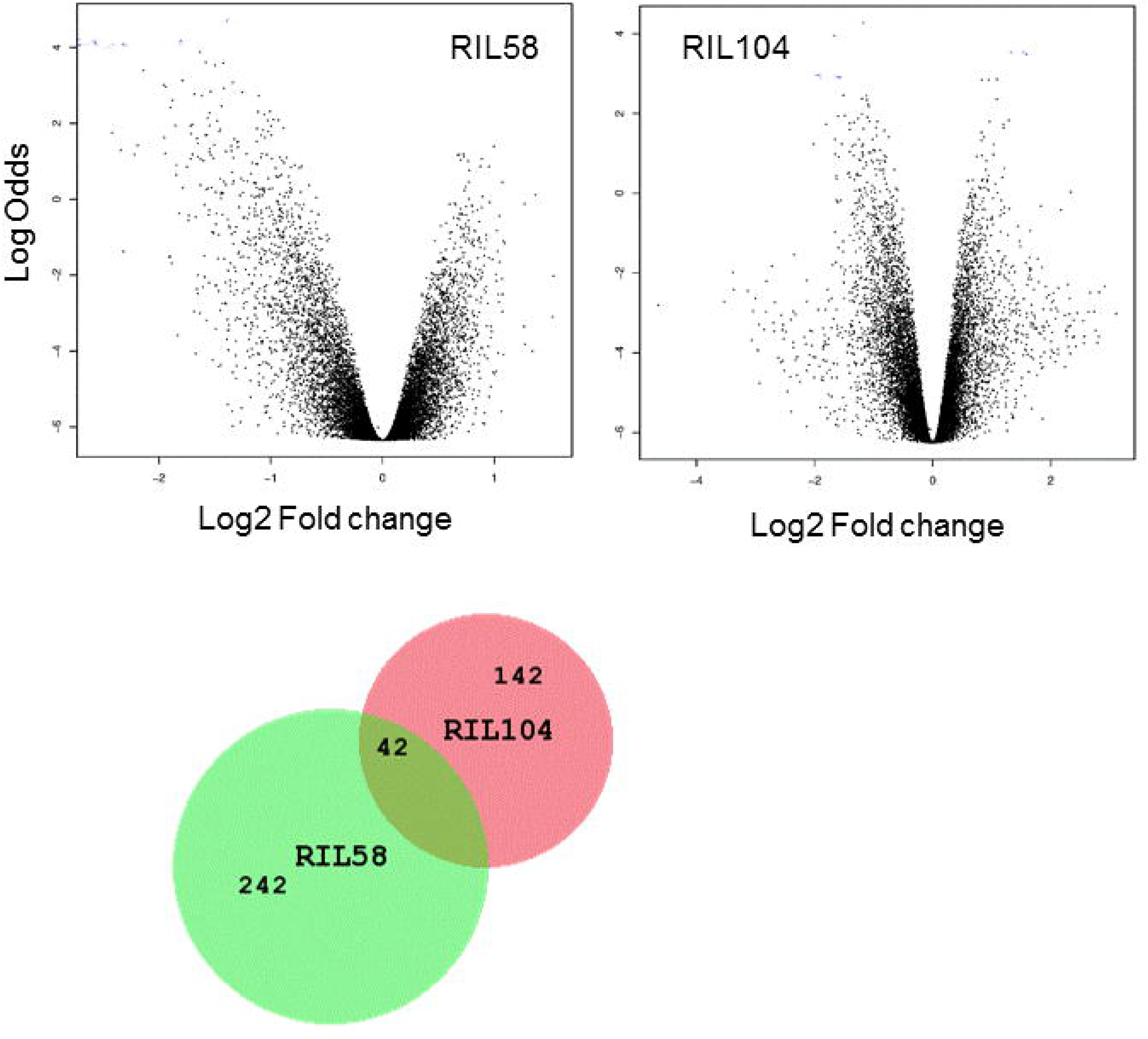
Differential expression induced by light in RI strains. Volcano plots of differential expression of RIL58 transcripts (left: 160 down-regulated and 25 up-regulated with light pulse) and RIL104 transcripts (right: 118 down-regulated and 56 up-regulated with light pulse). The empirical Bayes log-odds of differential expression are plotted against log2 fold change. Positive fold change represents up-regulation of transcript in the light - pulsed samples compared to the controls.

To identify eQTLs, we tested for genes that show significant difference in their expression between the two RILs strains (i.e. treatment: strain interaction; Table S7). Out of 131 transcripts that showed significant interaction (*p* < 0.001), four are located within QTL3 and could therefore may be considered as cis-QTLs. These were CG32189, Tsp74F (a membrane component), CG5577 (phosphatase and dephosphorylation activity), CG7510 (intracellular signal transduction and trans-membrane transport). All four loci are located within the *Df6411*deficiency that failed to complement in the complementation test.

We have also analysed the upstream sequences of the 131 candidate genes and searched for enrichment of regulatory motifs. There were 32 motifs that showed a significant enrichment (adjusted p-value p < 0.05, Supplementary Table S8). None of the 32 transcription factors is located within the deficiency intervals that failed to complement. Interestingly, one of the top enriched transcription factor motif Bab1 was previously associated with CLK/CYC targets (Meireles-Filho *et al*. 2014). Another interesting enriched motif is CF2 which is a transcription factor that regulates the microRNA mir-276a and has been recently shown to alter expression of *tim* in clock neurons (Chen and Rosbash. 2016).

## Discussion

Although light entrainment of the clock is a pathway that has been extensively studied, little is known about the genetic variation associated with its components. This type of information, which is addressed by QTL analysis is essential for understanding the evolution and the adaptive potential of this system.QTL studies of circadian photosensitivity were previously carried out in mammals. For example, Yoshimura and colleagues (Yoshimura *et al*. 2002b) crossed the retinally degenerated mice strains C57B*L*/6*J* and CB*A/J* (both homozygous for the mutation *rd*) that differed greatly in their phase response. They identified 5 QTLs on three different chromosomes, suggesting that multiple loci contribute to variation in light sensitivity, although no obvious candidate genes (e.g. photopigments) were mapped to these QTLs. Here, we have carried a QTL analysis of circadian photosensitivity in Drosophila, and identified a single major QTL in chromosome 2 (Fig. 2).

However, the evidence suggests that additional loci also contribute to photosensitivity variation. For instance, within the QTL2 interval, the 3133 deficiency that failed to complement the natural alleles does not span the *tim* locus, indicating that this large QTL interval may fractionate into multiple linked QTLs. In fact, the relatively small sample size used here (123 RILs) might have led to the Beavis effect (Beavis. 1998), where loci that are being failed to be detected due small power, contribute to size inflating and bias of the estimated proportion of the genetic variance explained by the nearest QTL.

Additional 3 QTLs were suggested by the multiple interval mapping (MIM) algorithm. The fact that the *cyc* locus, which is located within one of these QTLs (QTL3) was implicated in the complementation tests, provided further evidence that these additional QTLs may indeed contribute to the variation in light sensitivity.

The peak of the LOD of QTL2 coincides with the genomic position of *timeless,* and the *ls-tim* polymorphism that has already been shown to be involved in natural variation in light-sensitivity, and contributes to phenotypic variation that we observe here among the RIL. Besides demonstrating that the *ls-tim* SNP is a major component of the genetic architecture of this trait, the analysis of our RIL (from a population in Winters, California) indicates that this variation, which has emerged in Europe, is also segregating in North-American populations. Indeed, our recent survey of American populations (Pegoraro et al, submitted) shows that new ls-tim allele is present and follows a latitudinal cline, although this spatial pattern could reflect a demographical processes rather than natural selection, similar to the situation in Europe (Tauber *et al*. 2007).

QTL2 encompasses other loci worth noting and future research. *Lilliputian (lilli)* encodes a nuclear protein related to mammalian Fragile-X-Mental Retardation 2 protein FMR2 (Tang *et al*. 2001). The mammalian *fmr1*/*fmr2* has been shown to play a significant role in circadian function of the peripheral clock possibly through its defect in neuronal communication (Zhang *et al*. 2008). Similarly, Drosophila null mutations of the *Fmr1* are reported to alter circadian output with no detectable effect on the function of the central pacemaker (Inoue *et al.* 2002). Interestingly, Fmr1 was shown to be differentially expressed in response to light stimulation in a microarray study using Canton-S strain (Adewoye *et al.* 2015). In addition, LILLI has been shown to act as a dominant suppressor of activated MAPK pathway phenotypes (Dickson *et al.* 1996). The MAPK pathway has been reported to be activated by light, suggesting that its activation is vital for circadian light entrainment (Obrietan *et al.* 1998), and this signalling pathway has also been shown to mediate the function of PDF in *Drosophila* (Williams *et al*. 2001). Another gene of interest that is located within the QTL2 interval is *Shaw*, a member of the Shaker family encoding a voltage-gated K+ channels, which triggers hyperpolarisation of neurons upon activation. Misexpression of *Shaw* in clock neurons, particularly the dorsal neurons (DNs) results in disruption of the clock output and accumulation of PDF in projections of the ventral lateral neurons (LNv) (Hodge and Stanewsky. 2008). Interestingly, *Shaw* also showed a significant differential expression in response to light in both RIL58 and RIL104, although the similar transcriptional response in both loci rules out this gene as an expression QTL for this trait.

The substantial transcriptional variation revealed by the microarray experiments suggested that variation in many genes is associated with this phenotype (Fig 3). However, the overlap between the differential expression (total of 368 DEG) was rather modest (42 genes, 11%), which might be due to the rather limited power of the experiment (only two replicates per experiments). Alternatively, and not mutually exclusive, the lack of overlap suggests that the response to light evolves through different mechanisms that are highly dependent on the genetic background. This may also explain the little overlap with the light response of the Canton-S strain that has been previously studied (Adewoye *et al. 20*15). Only a single transcript, *sticks and stones (sns*), was shared between the RILs and the Canton-S lists (supp. Material). Interestingly, SNS was shown to be involved in the morphogenesis of the compound eye (Bao *et al.* 2010). The difference in the microarray platforms used here (spotted cDNA) and in the previous study (Affymetrix oligonucleotide microarrays) is likely to be an additional factor contributing to the differences between these studies.

Although *tim (and cry)* serve as a prominent light input for clock entrainment, it is clear that additional independent light inputs exist (Yoshii *et al*. 2015). These include three external photoreceptor organs, the compound eyes, the ocelli and the Hofbauer-Buchner eyelets. Clearly, each of these inputs may involve a plethora of proteins that might be targeted by natural selection. For example, it was shown that the compound eyes are essential for light entrainment under long days (Rieger *et al.* 2003), as well as for mediating moonlight response (Schlichting *et al*. 2014), a response that implicates light-sensing rhodopsins Rh1 and Rh6. A recent study (Hilbrant *et al.* 2014) demonstrated natural variation in rhodopsin expression (presumably reflecting variation in ommatidia patterns), which potentially may give rise to phenotypic variation in light entrainment of the clock. Although none of the rhodopsins were included in our DEG lists, one of the genes that show significant difference (i.e. RIL x treatment interaction) was *ninaA,* a gene that encodes cyclophilin, which transports opsins from the endoplasmic reticulum to the membrane. *ninaA* itself exhibits daily expression oscillation both in *Drosophila* (*Ueda et al*. 2002) and in two mosquito species (Rund *et al.* 2013).

Although the Drosophila PRC shows similar phase delay and advance responses (Li and Rosbash. 2013), the candidate genes we identified here for phase delay may be different from those involved in phase advance, reflecting different molecular mechanisms. For example, spectral composition of ambient light changes throughout the day, and was shown to be significant for light entrainment in birds (Pohl. 1999), fish (Pauers *et al.* 2012), and mammals (Walmsley *et al*. 2015). Thus, it is possible that during light pulse different photoreceptor populations are being activated in delay and advance zones respectively. Furthermore, it was suggested that the delay and advance responses are mediated by separate neural networks (Tang *et al.* 2010). A recent study using thermogenetics to activate different clock neuron clusters (Eck *et al.* 2016) showed that depolarization of the s-LNvs induced only phase advances, while stimulation of the LNds and the PDF-positive l-LNvs generates phase delays. Thus, variation in circadian photosensitivity may also be due to variation in the wiring of the clock neurons, which in turn might be due to genetic variation in developmental genes that drive the cellular connectivity of the clock.

Although expression analysis has been a popular way for identifying genes underlying QTLs, the approach has its own limitations (Verdugo *et al.* 2010). For instance, no significant gain is achieved by combining QTL/microarrays and more importantly, and often overlooked is the fact that transcriptional variation only captures a fraction of the genetic variation, as many genetic variations affect expression post-transcriptionally or post-translationally. The polymorphism in *timeless* is particularly an apt example, since the *ls/s tim* polymorphism is not manifested at the transcriptional level but rather translationally. In addition, we note that only a small fraction of DEGs that are induced by light may be in the QTL since a single transcription factor with a QTL may regulate many distant genes (i.e. trans eQTL).

Nevertheless, our study has generated a large number of candidate genes that may contribute to genetic variation in circadian photosensitivity, and the new genomic resources that have been recently made available, such as the Drosophila Synthetic Population Resource (DSPR) (Long *et al.* 2014), the Drosophila Genome Research Panel (DGRP) (Mackay *et al.* 2012) or the *Drosophila* Genome Nexus would allow finer mapping of variations associated with these genes.

## Data archiving

All microarray data used in this study have been deposited in GEO (http://www.ncbi.nlm.nih.gov/geo/) under accession number GSE77116.

## Acknowledgments

We would like to thank Dr Christopher Talbot and Dr Anne Genissel for their QTL mapping advise. We also thank the two anonymous reviewers whose comments and suggestions helped improve and clarify this manuscript. This work was funded by a BBSRC grant BB/G02085X/1 to ET.

